# Active and machine learning-based approaches to rapidly enhance microbial chemical production

**DOI:** 10.1101/2020.12.01.406439

**Authors:** Prashant Kumar, Paul A. Adamczyk, Xiaolin Zhang, Ramon Bonela Andrade, Philip A. Romero, Parameswaran Ramanathan, Jennifer L. Reed

## Abstract

In order to make renewable fuels and chemicals from microbes, new methods are required to engineer microbes more intelligently. Computational approaches, to engineer strains for enhanced chemical production typically rely on detailed mechanistic models (e.g., kinetic/stoichiometric models of metabolism) — requiring many experimental datasets for their parameterization—while experimental methods may require screening large mutant libraries to explore the design space for the few mutants with desired behaviors. To address these limitations, we developed an active and machine learning approach (ActiveOpt) to intelligently guide experiments to arrive at an optimal phenotype with minimal measured datasets. ActiveOpt was applied to two separate case studies to evaluate its potential to increase valine yields and neurosporene productivity in *Escherichia coli*. In both the cases, ActiveOpt identified the best performing strain in fewer experiments than the case studies used. This work demonstrates that machine and active learning approaches have the potential to greatly facilitate metabolic engineering efforts to rapidly achieve its objectives.

## INTRODUCTION AND BACKGROUND

In the near future, fuels and chemicals will have to be made renewably, and microbes are an attractive way to accomplish this due to their mild reaction conditions, product specificity, and product complexity. However, the number of commercial products made biologically is limited due to economic infeasibility and the incomplete understanding of biological systems resulting in numerous time-consuming iterations of the design-build-test cycle to optimize yields, titers, and/or productivities. While metabolic engineering aims to increase yield, titer, and/or productivities through genetic manipulations, it is often difficult to identify which genetic modification(s) (e.g., gene deletions, gene additions, and/or gene expression changes) are needed to improve biochemical production. To address this challenge, a variety of experimental and computational approaches have been developed in order to facilitate metabolic engineering efforts.

With a purely experimental approach, a large number of experiments may be needed to fully explore the potential genetic design space and find strategies that meet metabolic engineering objectives. Therefore, a number of high-throughput experimental approaches, including chemical genomics/BarSeq/TnSeq (that all quantify abundance of mutants in pooled libraries) *(1)(2)(3)*, MAGE (Multiplex Automated Genome Engineering) *(4)*, and TRMR (Trackable Multiplex Recombineering) *(5)* have been recently developed to improve metabolic engineering phenotypes, such as tolerance and chemical production. These experimental methods can rapidly generate large libraries of strains with high genetic diversity; however, these have only been applied to a relatively small number of microbial systems with metabolic engineering applications. Additionally, many of the techniques for identifying what genetic changes lead to desirable phenotypes rely on high-throughput screens or selections. Screening a large library of strains can be time consuming and requires a high-throughput method to monitor chemical production (e.g., colorimetric assays), which do not exist for many biochemicals, limiting the applicability of this approach. On the other hand, selections require a metabolic engineering objective connected to cellular growth or fitness. Such selections have been used to improve tolerance *(5)*, but it is more challenging to use them to find mutations that lead to greater metabolite production. Addressing these issues, experimental approaches such as multivariate modular metabolic engineering (MMME), which separates metabolic pathways into smaller modules that are varied simultaneously, can significantly reduce the design space to obviate the need for high-throughput screens. However, in doing so, valuable information is potentially lost and MMME still requires a semi-trial-and-error combinatorial construction of strains on the order of 10s, relying on human intelligence to deconvolute possibly complex, nonlinear interactions from sparse datasets to inform the next design *(6, 7)* Even so, most metabolic engineering projects still use a rational, iterative, trial-and-error approach that increases precursor and cofactor availability, alleviates bottlenecks, reduces flux through competing pathways, and expresses enzymes in biosynthesis pathways in order to increase desired production rate, product yield, or product titer.

Along with the experimental methods, a multitude of computational methods have been used to study microbial metabolic and/or regulatory networks and identify the genetic interventions needed to increase production of desired chemicals from low-cost substrates. These computational methods rely on mechanistic models (including genome-scale metabolic, kinetic, and regulatory models) or statistical models. Computational methods like OptKnock *(8)*, SimOptStrain *(9)*, and OptORF *(10)* rely on a stoichiometric, genome-scale, metabolic model to identify gene knockout and/or gene addition strategies that couple growth and metabolite production to enhance biochemical yields using experimental selections. Additionally, OptORF can also use integrated metabolic and transcriptional regulatory models to identify strategies involving metabolic and transcription factor gene knockouts and metabolic gene over-expression *(10)*. However, reconstructing a microbe’s transcriptional regulatory network is currently a major challenge and such integrated models exist only for well-studied organisms *(11)(12)(13)*. Alternatively, kinetic models, which are much more detailed than stoichiometric metabolic models, can be used to increase flux through a pathway *(14)(15)(16)(17)(18)*. However, due to the complexity of biological systems and incomplete datasets, there is much uncertainty attached to parameters within kinetic models. To address this, computational workflows such as ORACLE and iSCHRUNK are being developed that utilize kinetic models, metabolic control analysis, and machine learning principles to minimize kinetic parameter uncertainty to suggest engineering strategies in the absence of complete information *(19, 20)*. Nevertheless, these kinetic models require costly, time-consuming, and complex datasets (e.g., fluxomic, proteomic, and metabolomic), as well as a thorough understanding of substrate-level regulation, to accurately parameterize them, limiting kinetic modeling to well-studied organisms.

In contrast to mechanistic models, which often require large datasets to build them, statistical models can be used instead. Design-of-experiments tools, such as JMP *(21)* and DoubleDutch *(22)*, can be used to design an initial set of experiments that evaluate the impacts of genetic mutations on desired metabolic engineering objectives. However, design-of-experiments tools often lack capabilities to use these initial experimental results to design the next set of experiments. Recently, machine learning approaches have been used to optimize gene expression levels to enhance metabolic flux through desired pathways. Lee and colleagues used a categorical log-linear regression model to predict how different promoters, used to drive expression of biosynthetic genes, impacted violacein titers *(23)*. Farasat et al., in addition to their mechanistic kinetic model, used non-mechanistic models (i.e., a geometric and two statistical linear regression models) to predict how different ribosome binding sites (RBSs), controlling expression of three different biosynthesis genes, affected neurosporene *(14)*. While these non-mechanistic models could accurately predict the performance for new combinations of previously tested RBSs or promoters (referred to as exploration), they were unable to predict the performance of gene expression constructs containing new RBSs or promoters (referred to as extrapolation).

Here, we developed an active and machine learning-based approach to design gene expression constructs for metabolic engineering—ActiveOpt—that overcomes many of the aforementioned drawbacks. Although this is the first reported study that uses active learning-in metabolic engineering, active learning has been previously used in a wide range of other applications *(24),(25),(26),(27),(28),(29),(30)*. ActiveOpt integrates computational and experimental efforts to improve metabolic engineering objectives using substantially fewer and simpler experiments (e.g., measuring biochemical yield or productivity) than many state-of-the-art approaches. ActiveOpt combines active and machine learning techniques without the need for detailed mechanistic models of the underlying metabolic and regulatory networks or a large initial experimental dataset. ActiveOpt guides the search for effective genetic engineering strategies using a machine learning classifier with simple inputs (e.g., predicted RBS strengths) constructed from at least two experimental results. As more results from new experiments become available, a classifier is refined to improve the selection of the next set of experiments. This cycle between classifier refinement, biochemical yield or productivity prediction, and experimental testing stops when either the metabolic engineering objective stops improving substantially, or a maximum number of experiments has been performed.

In this study, we show how ActiveOpt identified optimal combinations of genes and RBSs needed to increase biochemical yields or productivities for two different metabolic engineering case studies. Specifically, in the two case studies, we show that a simple machine learning classifier can accurately make qualitative predictions of product yield (i.e., low or high yield) from gene choices and RBS strength predictions *(31),(32)* using very few experiments, without requiring a detailed mechanistic model. Second, we show that ActiveOpt identifies combinations of RBSs and genes with the highest valine yields and neurosporene productivities in fewer experiments than a random trial-and-error approach. Third, four additional combinations of gene expression constructs predicted by ActiveOpt to have high valine yields were experimentally verified after prediction from ActiveOpt. Finally, we show that ActiveOpt can be used to predict the outcomes of both exploration and extrapolation experiments, indicating that new combinations of previously tested and un-tested gene expression constructs can be selected in the experimental design process. Together, these results show the potential effectiveness of using ActiveOpt for metabolic engineering applications.

## RESULTS

An active learning and machine learning approach (ActiveOpt) for designing experiments was developed and applied to two metabolic engineering cases studies, one of which is reported for the first time here. We evaluated the accuracy of a machine learning classifier to predict valine yields from RBS strength estimates—the same classifier used by ActiveOpt. Although most of the experimental dataset for this case study was generated without using ActiveOpt, no knowledge of the experiments or valine production except for the pathway was used to evaluate ActiveOpt’s performance. ActiveOpt’s performance at identifying the genetic parts that maximize yield or productivity in the fewest possible experiments was evaluated using three different methods for selecting experiments. Four new combinations of previously tested RBSs (i.e., exploration experiments) were suggested by ActiveOpt and tested experimentally; experimental results for these four new combinations were not available when ActiveOpt was used to make the prediction. Similarly, ActiveOpt was applied to enhance neurosporene productivity in *E. coli* using data from previously published experiments *(14)*, and RBSs not used during ActiveOpt training (i.e., extrapolation experiments) were selected to improve neurosporene productivity.

### Metabolic Engineering of *E. coli* for Valine Production

Valine is an amino acid widely used as a nutritional supplement in several industries with a demand of about 500 tons annually *(33)*. Amongst engineered *E. coli* valine production strains, the highest reported elemental carbon yield is 39% supplied C converted to valine *(34)*; however, the strain requires supplementation with yeast extract, acetate, leucine, isoleucine, and D-pantothenate. Our goal was to engineer an *E. coli* strain with higher valine yields but without complex media requirements. Plasmids expressing valine biosynthesis and exporter genes (either *ilvBN*DE, ilvIH*C-ygaZH*, or *ilvIH*C*--ygaZH*, Figure 1) were designed using rational approaches, such as performing carbon balances to identify bottlenecks, using engineered enzymes, and identifying trends and testing systems-level hypotheses based on collected data. However, computational approaches were not used to design experiments. The two plasmid backbones, promoters, gene number, and order were fixed throughout the study with variations allowed for one gene *(ilvC* or *ilvC*)* and individual enzyme RBS strengths. A total of 39 plasmids were constructed and tested in 89 pairwise combinations before the best strain was identified which achieved an elemental carbon yield of 45% (or 54.7% of the maximum theoretical (MT) yield from glucose and acetate) in a defined minimal medium—the highest carbon yield reported in *E. coli*(Figure 2A). A total of 49 pairwise combinations were tested before one of the top strains (reaching ~90% of the best strains % of MTY); see supplementary information for details on the strategy employed for all 89 experiments.

**Figure 1:**
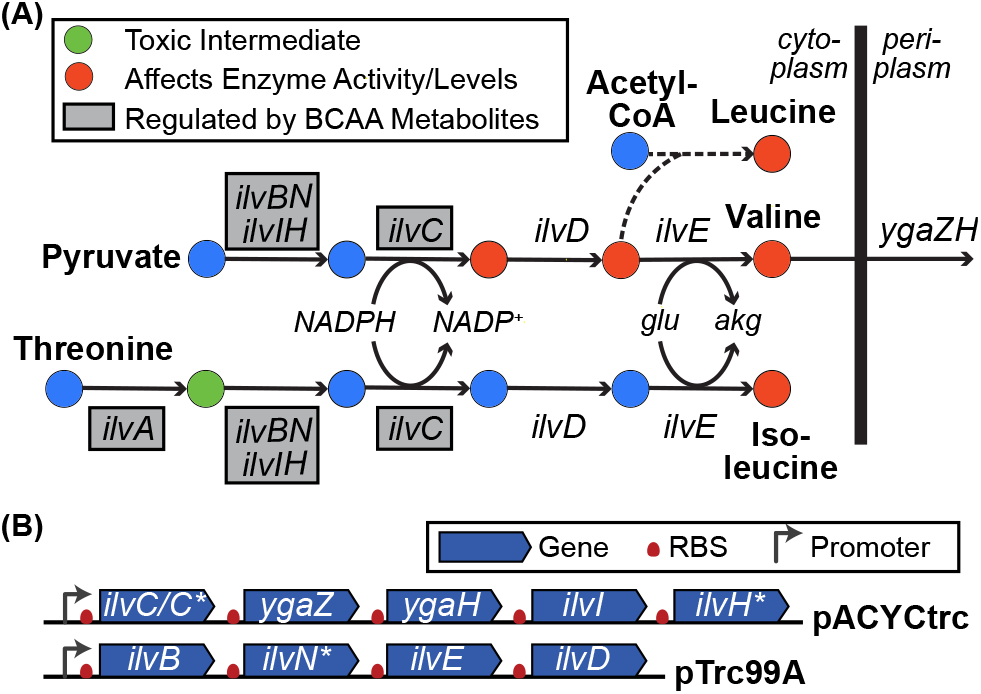
Biosynthesis pathway for branched chain amino acids in *E. coli*. There are nine genes involved in valine export and biosynthesis from pyruvate. The dashed arrow indicates the need of multiple reactions to convert acetyl-CoA and 3-methyl-2-oxobutanoate to leucine. Metabolites that regulate branched chain amino acid biosynthesis enzyme activity or levels are shown in red. Metabolites that are toxic are shown in green. Enzymes that are regulated by branched chain amino acid metabolites are boxed in grey.

**Figure 2:**
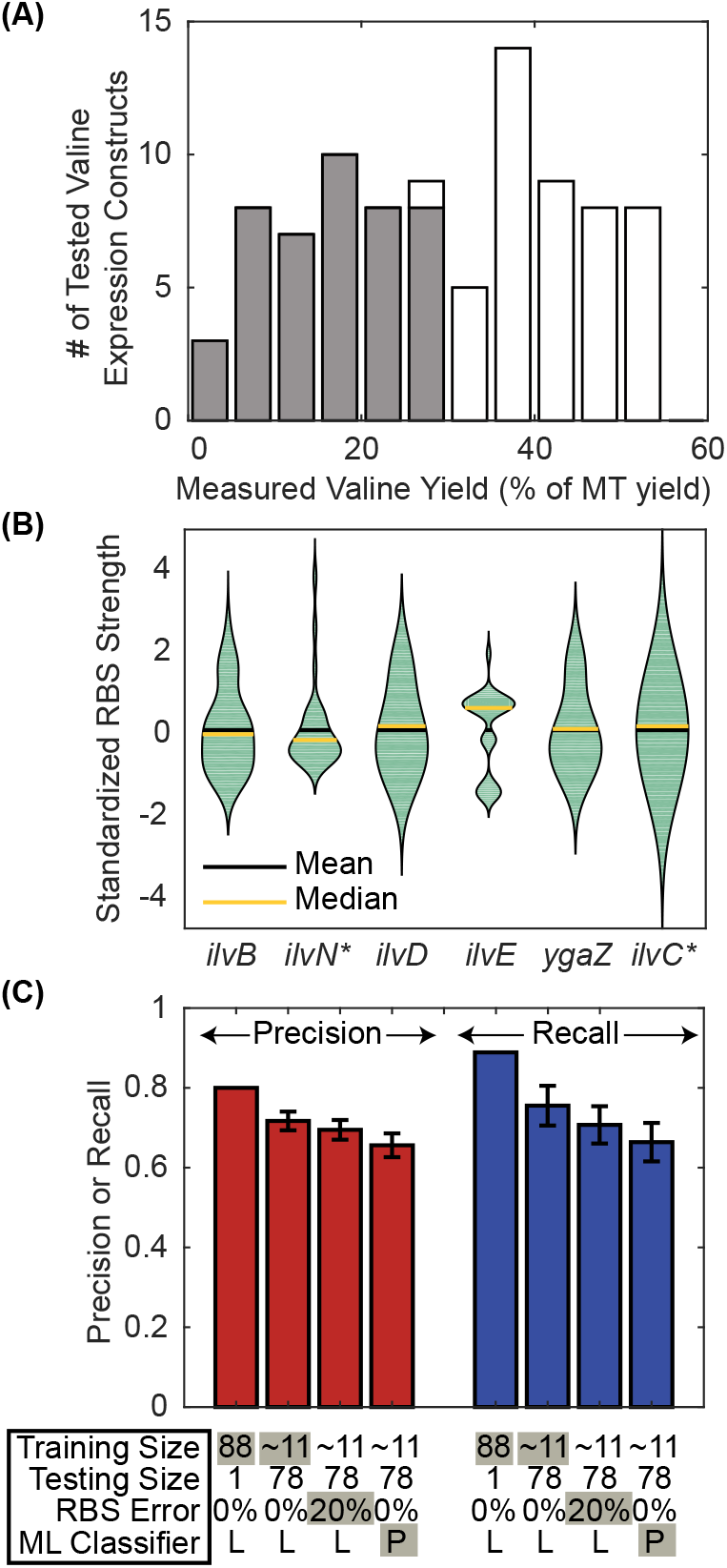
Machine learning approaches applied to the valine experimental dataset. Panel (**A**) shows a histogram of the valine yield in all 89 experiments and whether they were classified as high (white bars, 46 experiments) or low (grey bars, 45 experiments) yield. (**B**) Shows a violin plot (where the outer shape width is proportional to frequency of occurrence and the black and yellow bars indicates the mean and median values, respectively) of the standardized RBS strengths (see Methods for details) for each gene whose RBS varied across the experiments. The precision and recall are shown in panel (**C**) for four different cases with different training (and testing) set sizes, added RBS strength errors, and with linear (Lin.) or non-linear (Non-Lin.) classifiers. Precision (red bars) is the ratio of true positives (i.e., correctly predicted high yield experiments) to the total predicted positives (i.e., total predicted high yield experiments), whereas, recall (blue bars) is the ratio of true positives to the total actual positives (i.e., total actual high yield experiments). The bar represents the average and the error bars show the standard deviation across 1,000 inverse eight-fold cross-validations.

### Machine Learning Algorithms Accurately Predict Valine Yields

A total of 89 different valine production experiments were used to evaluate how well different machine learning classifiers could qualitatively predict valine yields (i.e., high or low yield) from RBS strengths and enzyme choices. All valine experiments were classified as either high yield (45 experiments) or low yield (44 experiments) using a fixed cutoff of 29% of the MT yield of valine from glucose and acetate, so that a randomly chosen experiment has roughly a 50% chance of being high yield (Figure 2A). The input data used by the machine learning classifiers included the RBS strength predictions for 6 of the plasmid-expressed genes (i.e., the genes whose RBSs were varied across experiments, Figure 2B) and whether a native *ilvC* or mutated *ilvC* (35)* was used (encoding the NADPH and NADH-dependent enzymes, respectively). The resulting classifier’s qualitative output was either a high or low valine yield prediction for a given experiment from a set of inputs.

To determine first if a linear Support Vector Machine (SVM) classifier *(36)* could accurately predict a valine experimental outcome correctly, we performed a leave-one-out crossvalidation (LOOCV). In this case, the results from 88 experiments were used to train an SVM classifier and the classifier was used to predict the final experimental outcome. This was repeated 89 times, with each experiment being left out of the initial training dataset used to build the classifier. The precision (the fraction of experiments that were predicted to be high yield which were found to have high yields experimentally) and recall (the fraction of high yield experiments that were predicted to be high yield) were calculated from this LOOCV analysis and are shown in Figure 2C. The precision and recall was 0.80 and 0.89, respectively, across these 89 different linear SVM classifiers. The agreement between machine learning model predictions and experimental outcomes was statistically significant (p-value = 1.35×10^−10^ using a Fisher Exact Test).

Given the high level of accuracy for the linear classifiers, additional analyses were performed to evaluate whether fewer experiments could be used to train the classifier, if errors in predicted RBS strengths would impact accuracy, and if non-linear classifiers could improve predictions. In each case, the 89 possible experiments were randomly assigned to one of eight folds (or groups), with each fold including ~11 experiments. Each fold was used independently as a training set to build a classifier, which was used to predict the outcomes for experiments in the seven other folds. The precision and recall values were calculated using predictions from all eight independent classifiers. This inverse eight-fold cross-validation was then repeated 1,000 different times and the resulting precision and recall values were averaged. When the number of experiments used to train the classifiers was lowered from 88 to ~11, the average precision (0.72) and recall (0.76) across 1000 inverse eight-fold cross-validations reduced only slightly (Figure 2C). Additional fold sizes were also investigated, containing between ~5 and ~45 experiments, with precision ranging between 0.67 and 0.79 and recall ranging between 0.68 and 0.87 (Supplementary Figure S1). Since the RBS Calculator *(31)* used to calculate the translation initiation rate may be inaccurate, it could potentially produce erroneous classifier input data. To evaluate the impact of potential errors in RBS strength predictions, the calculated RBS strength *(31)* was randomly changed up to +/− 20% for each of the 6 genes whose RBS sequence was varied. Once again, 1000 inverse eight-fold cross-validations were generated (by randomly assigning ~11 experiments to one of eight folds) and the precision and recall were calculated across all eight folds. From this analysis, 20% errors in the predicted RBS strengths by the RBS calculator did not significantly affect the precision and recall rates (Figure 2C). Finally, a non-linear polynomial classifier was tested to see if it could improve machine learning model predictions, but the results were similar to the linear classifier with an average precision of 0.66 and recall of 0.66 (Figure 2C). While precision and recall were not found to be very sensitive to fold-size, RBS errors, or classifier type, the precision and recall were sensitive to the cutoff used to classify experiments as high/low yield. In this case, the precision and recall of the classifier decreased as the fraction of experiments that were classified as high yield decreased (Supplementary Figure S1), since there are fewer high yield cases to learn from. Hence, we proceeded to use a linear SVM classifier, with a cutoff that results in proportionately high and low yield cases, and without any RBS strength errors for all subsequent analyses.

### Comparison of Different Active Learning Approaches

In total, 89 valine experiments were performed initially; however, if the study was repeated, could we identify the highest yielding strains in fewer, more intelligently selected experiments? To answer this, two active learning algorithms—ActiveOpt and Upper Confidence Bound *(37)* (UCB) — were applied to maximize valine yields in fewer experiments. For ActiveOpt, a small number of starting experiments (e.g., 2 or 3) were selected (Figure 3B) and an initial high/low yield cutoff was calculated (equal to the average of the highest and lowest yield across the set of selected experiments). Results from these experiments were used to train an initial linear SVM classifier (in the case of ActiveOpt) or a Gaussian process regression model (in the case of UCB). To identify the “next experiment” to be conducted and added to the training set used to generate subsequent classifiers and yield cutoffs (Figure 3A), we investigated three approaches with ActiveOpt (referred to as next-experiment selection approaches):

1. Closest-to-the-Hyperplane: with this approach, the closest experiment to the SVM hyperplane that is predicted to be high yield and has not been performed yet is chosen. This active learning approach could potentially generate accurate classifiers more quickly because experiments with the most uncertainty in their outcome (since they are close to the SVM hyperplane) are performed first.
2. Farthest-from-the-Hyperplane: with this approach, the farthest experiment from the hyperplane that is predicted to be high yield and has not been performed yet is chosen. This active learning approach could potentially reach the highest yielding strains in the fewest number of experiments.
3. Farthest-then-Closest-to-the-Hyperplane: with this approach each next experiment alternates between either being farthest from the classifier’s hyperplane or closest to the hyperplane on the high yield side. This active learning approach could attempt to achieve two objectives: reaching the highest yielding strains and building an accurate classifier.

**Figure 3:**
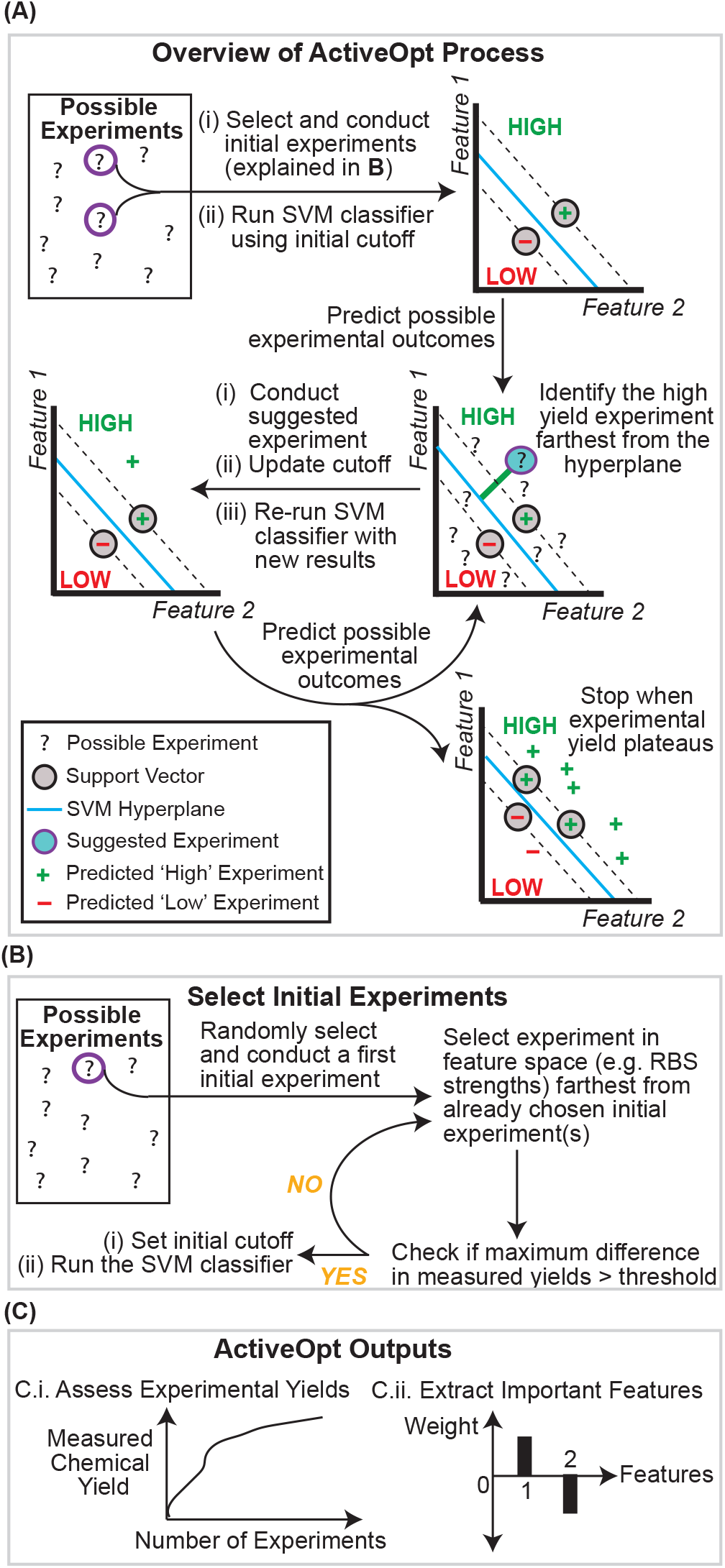
Overview and Output of ActiveOpt. Panel (**A**) shows a Flowchart of the **ActiveOpt** method. Panel (**B**) shows the process for selecting the initial set of experiments on which the classifier is initially run. Panel (**C**) shows possible outputs generated by **ActiveOpt**, such as: maximum product yield found versus number of experiments performed or identification of important features affecting product yield.

We then compared ActiveOpt and UCB performances to a random trial-and-error approach (where the next experiment was randomly chosen from the set of remaining unperformed experiments). While ActiveOpt (Figure 3) and UCB are active learning algorithms, the random selection approach is not an active learning approach since current information is not used to inform selection of the next experiment.

To avoid biasing the comparisons by only selecting a single initial experiment, we ran the random scenario 1000 times, where each time an initial experiment was randomly chosen and then each of the 88 remaining experiments were randomly selected one by one. ActiveOpt was run with each of the 89 experiments used as the initial experiment for each of the three next-experiment selection approaches described above. At each iteration, ActiveOpt used the updated linear SVM classifiers from the previous round of data to select the next experiment (Figure 3A). ActiveOpt selected experiments to perform until no unperformed experiments were predicted by the SVM classifier to be high yield (i.e., all remaining potential experiments were predicted to be low yield). For the random selection approach, another experiment was performed until no additional experiments were available from the set of 89 experiments.

For each run, we first determined how many total experiments had to be performed before a satisfactory strain was found that had at least 95% of the highest observed valine yield across all 89 experiments (the highest observed elemental carbon yield was 45%, which is 54.7% of the MT yield). Figure 4A–C shows histograms of the total experiments needed to find a satisfactory strain across the ActiveOpt runs using different next-experiment selection approaches (see Supplementary Figure S2 for farthest-then-closest-to-the-hyperplane results). It is possible to identify that the farthest-from-the-hyperplane approach frequently finds a satisfactory valine production strain in fewer experiments than the other approaches (although farthest-then-closest-to-the-hyperplane and closest-to-the-hyperplane approaches are still an improvement over random sampling, a non-active learning approach). In 59 out of 89 cases, fewer than 10 expression constructs had to be tested before a satisfactory strain was found using the farthest-from-the-hyperplane approach compared to 475 out of 1000 or 41 out of 89 cases for the randomly chosen or closest-to-the-hyperplane approaches, respectively (Supplementary Table S3). This result shows that an active learning approach (where continually updated information is used to design the next experiment) can reduce the amount of time and effort needed to generate high yield strains.

**Figure 4:**
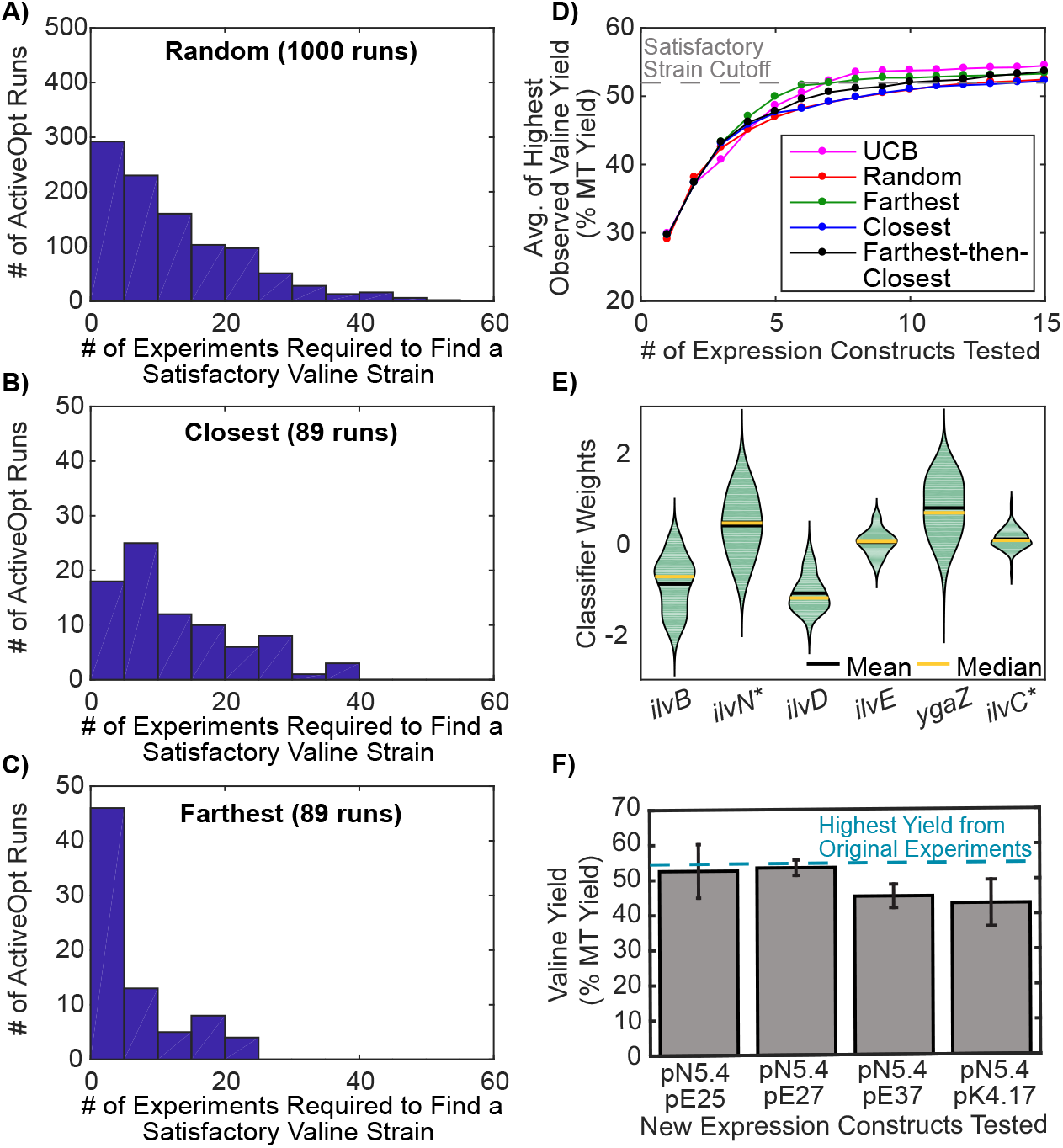
ActiveOpt Applied to Enhance Valine Yield. Panels (**A-C**) show histograms for the number of total experiments needed by ActiveOpt to identify a satisfactory strain (i.e., a strain with a yield >95% of the highest observed valine yield across all experiments) using different “next experiment” selection approaches when 89 different first initial experiments were used to start the algorithm. Panel (**A**) used random selection. Closest-to-the-hyperplane was used in panel (**B)**, and farthest-from-the-hyperplane in panel (**C**). In panel (**D**), the average from the 89 ActiveOpt or UCB runs of the highest observed % valine yield is plotted as a function of the number of total experiments performed. Panel (**E**) shows the distribution (using violin plots where the outer shape width is proportional to frequency of occurrence and the bar indicates the average value) of the feature weights from the final classifiers generated from the 89 ActiveOpt runs using the farthest-from-the-hyperplane experimental selection approach. An SVM classifier was built from the original 89 experiments and used by ActiveOpt to identify four new experiments (not included in the original 89 experiments) that were farthest from the classifier’s hyperplane. In all four new experiments the valine yields were high (panel **F**) as predicted by ActiveOpt.

Another way to evaluate the performance of the different approaches is to identify, at each iteration (i.e., new experiment selection), the highest observed yield across the subset of currently performed experiments. This highest observed yield can then be averaged across the 89 runs with different starting experiments. From Figure 4D, it can be seen that the farthest-from-the-hyperplane approach steeply increases the valine yield per experiment, as compared with other next-experiment selection approaches. The slope of the plot in Figure 4D can also be used as an indicator to decide whether to perform more experiments or not (e.g., after 7 experiments the curve plateaus for the farthest-from-the-hyperplane approach). The final classifiers (when no more experiments were predicted to be high yield) at the end of each of the 89 ActiveOpt runs were more accurate when closest-to-the-hyperplane approach was used (with average precision and recall of 0.91 and 0.69 for all 89 experiments, respectively; and with standard deviations for precision and recall of 0.03 and 0.18, respectively), compared to the other next-experiment approaches (Supplementary Table S3).

In addition to using ActiveOpt with an SVM classifier, the UCB active and machine learning algorithm was evaluated, which allows tradeoffs between exploration and exploitation *(37)*. UCB uses a regression model’s predictions and confidence intervals to maximize an unknown function, in this case valine yield. Here, UCB used a Gaussian process regression model to predict valine yields, as compared to the SVM classifier used by ActiveOpt, which predicts high/low yield. Both UCB and ActiveOpt, on average, would take 8 experiments to find a satisfactory strain (Figure 2D). For a small number of valine experiments (between 3 and 6) ActiveOpt performs slightly better than UCB, while UCB performs slightly better than ActiveOpt after 8 experiments (Figure 2D). These results show that ActiveOpt and UCB can very accurately and efficiently identify high yield strains using results from a small number of experiments (e.g., ~8 in the valine case), nearly an order of magnitude less than the total 89 experiments originally performed to achieve the same yield.

### Significant Features from Resulting Machine Learning Classifiers

Machine learning classifiers can also be used to identify feature weights, a relative measure of the sensitivity of the linear SVM classifier output (in this case yield) to changes in feature value inputs (e.g., RBS strengths). Figure 4E shows the distribution of weights for the final classifiers (i.e., when no more high yield experiments are predicted for each of the 89 runs with unique initial experiments) when the farthest-from-the-hyperplane approach is used by ActiveOpt. From Figure 4E, it can be seen that *ilvB* and *ilvD* have strong negative weights in most of the runs, while *ilvC*, ilvN\** and *ygaZ* have positive weights. Increasing the RBS strengths of the genes with positive weights and decreasing the RBS strengths of the genes with negative weights should result in strains with high valine yields. Multinomial logistic regression (which fits binary outcomes to continuous input features) was also used to compare features from the valine dataset (Table 1). It can be seen that only the coefficients for *ilvB, ilvN*, ilvD* were significant, with a p-value less than 0.05. However, the signs of the weights were similar to those predicted by ActiveOpt, further supporting the utility of the machine learning approach.

**Table 1:** Feature weights from Logistic Regression, ActiveOpt (using farthest-from-the-hyperplane approach), and UCB.

| **RBS**<br>**Strength**<br>**for Gene** | **Logistic Regression**<br>**Coefficients**<br>**(p-values)** | **Average**<br>**ActiveOpt**<br>**(w/Furthest)W**<br>**eights** |
| --- | --- | --- |
| *ilvB* | -2.38 (0.017) | -0.90 |
| *ilvN\** | 3.50 (0.023) | 0.42 |
| *ilvD* | -4.03(0.001) | -1.10 |
| *ilvE* | 0.43 (0.490) | 0.02 |
| *ygaZ* | 0.16 (0.700) | 0.77 |
| *ilvC\** | 0.27 (0.545) | 0.09 |

### Newly Designed Valine Experiments by ActiveOpt

ActiveOpt suggested four new exploration experiments, using new plasmid combinations of previously tested RBSs, which were farthest from the hyperplane using a linear SVM classifier trained on all 89 previous experiments. Figure 4F shows that the valine yields in all four new experiments were correctly predicted to be high yield (>=29% MT yield), with one combination being 53.4% MT yield, very close to the highest yield (54.7% MT yield) from the original 89 experiments. Therefore, if distance from the hyperplane is indicative of valine yield, then no additional experiments, using combinations of existing plasmids (exploration), are predicted by ActiveOpt to increase yields above those found in the 93 experiments performed. Similarly, UCB predicted no untested plasmid pair combinations would have greater valine yields than those already tested.

### Application of ActiveOpt to Enhance Neurosporene Production

Farasat et al. *(14)* recently reported a neurosporene productivity dataset in *E. coli* that used a designed RBS sequence library to vary expression of three neurosporene biosynthesis pathway genes *(crtEBI)* (Figure 5A). The authors initially designed 73 expression constructs for *crtEBI*,transformed them into *E. coli*, and measured the specific neurosporene productivity (exploration experiments, Figure 5B). Next, a kinetic model (capable of extrapolating designs) was built for the 24 elementary reactions in the neurosporene biosynthesis pathway to design 28 new expression constructs (extrapolation experiments), increasing neurosporene productivity from 196.3 to a maximum of 286 μg/gCDW/hr.

**Figure 5:**
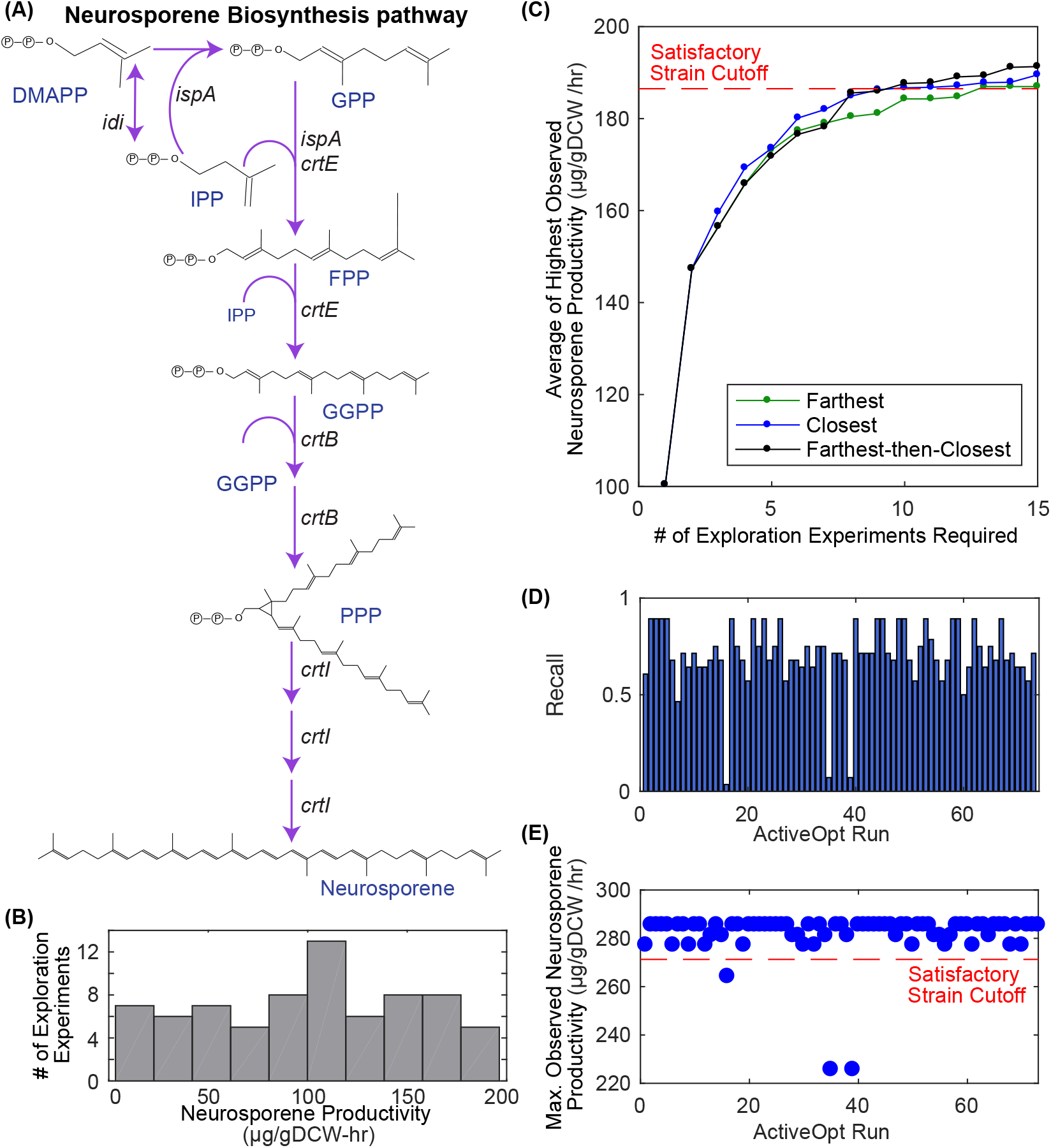
ActiveOpt Applied to Enhance Neurosporene Productivity. Panel **(A)** shows the neurosporene biosynthesis pathway. Panel **(B)** shows the neurosporene productivity measured by Farasat et al. in the exploration experiments. Panel (**C**) shows the average of the maximum observed neurosporene productivity found across the 73 ActiveOpt runs using different approaches for finding the next experiment (farthest-from-the-hyperplane = green, closest-to-the-hyperplane = blue, and farthest-then-closest-to-the-hyperplane = black). Panel (**D**) shows for each of the 73 final extrapolation ActiveOpt cutoffs and classifiers (using first exploration then extrapolation experiments) what the recall was for the extrapolation experiments (using new RBSs not tested in the exploration experiments). Panel (**E**) shows for each ActiveOpt run (using first exploration then extrapolation experiments) with the farthest-from-the-hyperplane approach what the maximum observed neurosporene productivity would have been across selected extrapolation experiments.

This initial exploration dataset was used by ActiveOpt to test whether the most productive strains could be identified in fewer than 73 experiments. Figure 5C shows the average highest observed neurosporene productivity as a function of the chosen number of exploration experiments for several next-experiment ActiveOpt approaches. In this case, ActiveOpt was run with each of the 73 exploration experiments performed by Farasat et al. as the initial experiment. This figure also indicates that ActiveOpt identified strains with at least 95% of the best productivity from the exploration experiments in much fewer experiments than the 73 experiments performed by Farasat and colleagues. On average, a satisfactory strain (with a productivity of >186.5 μg/gCDW/hr) would have been found with ~10 experiments for the closest-to-the-hyperplane and farthest-then-closest-to-the-hyperplane approaches and ~13 experiments for the farthest-from-the-hyperplane approach (Supplementary Table S4). Notably, ActiveOpt does not require any kinetic information to optimize expression constructs for the biosynthesis pathway. Furthermore, Farasat and colleagues found that high neurosporene productivity requires high *crtE* activity, agreeing with the final average ActiveOpt classifier weights of 1.07, −0.03, and 0.09 for *crtE, crtB*, and *crtI*, respectively, for the farthest-from-the-hyperplane approach (Supplementary Figure S3 and Supplementary Table S5).

The first 73 exploration experiments performed by Farasat et al. explored the design space for RBSs controlling neurosporene production. Using a kinetic model, the authors designed new RBSs predicted to further increase neurosporene production resulting in 28 new extrapolation experiments (since the RBSs were previously untested). The 73 final ActiveOpt classifiers (when no more high productivity exploration experiments were predicted) generated from the exploration experiments were each used to choose an extrapolation experiment with the farthest-from-the-hyperplane approach. ActiveOpt was then allowed to continue selecting new extrapolation experiments, by updating the cutoff and classifier, until no remaining extrapolation experiments were predicted by ActiveOpt to have high productivity. The final recall for the extrapolation experiments across all 73 runs (when ActiveOpt was started with final classifiers from the exploration experiments) had an average of 0.70 and standard deviation of 0.17 (Figure 5D and Supplementary Table S4). Of the 73 ActiveOpt runs, 47 would have found the highest productivity extrapolation experiment (286 μg/gCDW/hr), 58 would have found one of the top two productivities, and 70 would have found a satisfactory strain with >271 μg/gCDW/hr neurosporene productivity (Figure 5E). Slightly more runs identified a satisfactory strain when the closest-to-the-hyperplane and farthest-then-closest-to-the-hyperplane approaches were used with ActiveOpt (Supplementary Table S4). The average number of extrapolation experiments needed to find a satisfactory strain was 2, 4, and 6 when closest-to-the-hyperplane, farthest-then-closest-to-the-hyperplane, and farthest-from-the-hyperplane approaches were used, respectively (Supplementary Figure S4). This is substantially less than the total 28 extrapolation experiments performed by Farasat and colleagues. Together, these results show that ActiveOpt can be applied to extrapolation experiments involving previously untested RBSs.

## DISCUSSION

Machine learning uses statistical models to identify non-intuitive patterns between input features and experimental outcomes and has been applied to a wide range of fields; however, its use in metabolic engineering has been limited. We evaluated whether machine learning could be used in an active learning framework (ActiveOpt) to accelerate development of biochemical production strains. ActiveOpt was applied to two separate datasets, a published dataset for neurosporene productivity and a new valine dataset reported here—the latter of which achieved the highest reported *E. coli* valine yield in a defined minimal medium. We showed that a linear classifier is able to qualitatively predict yields with high precision and recall using only predicted RBS strengths and gene choices *(ilvC* or *ilvC*)* as inputs. When this machine learning classifier was integrated into an active learning framework, satisfactory strains could be identified in significantly fewer design iterations than the original experimental studies. In particular, there does not seem to be a need for a non-linear classifier.

ActiveOpt is a method for efficiently exploring the design space to identify the subset of gene expression constructs which give rise to strains with higher yields or productivities. Since ActiveOpt does not rely on high-throughput selections or screens to identify these optimal expression constructs, this approach could be applied to enhance production of a larger number of biochemical targets. ActiveOpt has low upfront requirements, in terms of data and understanding of the metabolic pathway, only requiring predicted RBS strengths and measured yields/productivities. Since ActiveOpt does not rely on detailed mechanistic or kinetic models it does not require large, complex ‘omics datasets to parameterize them. An important advantage of ActiveOpt, relative to most other supervised machine learning applications, is its ability to predict experimental outcomes outside the training set design space (i.e., extrapolation experiments) to achieve better results.

ActiveOpt also identifies the features that most significantly affect the metabolic engineering objective (in our case RBS strengths), which might be useful in further shrinking the design space for future studies on a similar pathway or narrowing the focus of the current study. Feature selection can direct our attention to portions of the pathways where a more detailed model or mechanistic insights into the system might be necessary to fine tune yields/productivities. Analysis of these features was useful in both case studies, and in the neurosporene study the feature weights for the genes found by ActiveOpt were consistent with conclusions drawn from a more detailed kinetic model of the pathway.

This work shows how machine and active learning can be used to successfully streamline the development of high biochemical production strains. While machine learning models worked well for the two case studies evaluated in this work, it is possible that optimizing flux through other metabolic pathways might require other types of classifiers and/or regressors to achieve accurate predictions. Future work should evaluate ActiveOpt’s performance on other metabolic engineering targets and investigate whether design decisions can include other types of gene expression control elements (e.g., promotors and terminators). The performance and validation of ActiveOpt opens avenues for its implementation to guide projects with a defined parameter design space from inception to outcome. While not explicitly tested here, this would be a true test for method robustness and would validate machine learning algorithms as a useful tool for metabolic engineers.

## METHODS

### ActiveOpt: Active Learning using a SVM classifier

ActiveOpt uses a SVM classifier *(36)* to perform active learning *(25)*. The built-in MATLAB SVM classifier function (‘svmtrain’) was used for binary classification (“high” and “low”) of biochemical yield or productivity data obtained from experiments. For both the valine and neurosporene cases the predicted RBS strengths for the individual genes in the biosynthesis pathways were used as features for classification and the set of all possible RBS strength values defines the feature space. For the valine dataset, if a gene was not included on a plasmid (i.e., *ilvC* or *ilvC*\*) then the associated RBS strength was set to zero. The predicted RBS strengths (from the RBS Calculator *(31)*) were standardized for each gene by subtracting the mean RBS strength and dividing by the standard deviation across all the values in the design space.

A machine learning classifier finds a decision boundary, a hyperplane in the multidimensional feature space, to predict whether a collection of feature values would result in either “high” or “low” yield/productivity. The linear SVM classifier requires experiments from each group be included in the training set. In the event that a fold was created that included experiments from only one group, then data from all other assigned folds were excluded from the analysis and the MATLAB ‘crossvalind’ function was used again to randomly assign all experiments to the specified number of folds. This random process was repeated for the inverse fold cross-validation until 1,000 appropriately assigned folds were found (i.e., each fold has both and high and low yield experiments).

ActiveOpt needs few starting data points to train the initial classifier and then ActiveOpt predicts all other experimental outcomes. For the initial set of experiments, ActiveOpt selects one experiment and then chooses another initial experiment from the available experiments which has maximum Euclidean distance in the feature space from the first chosen experiment (Figure 3A-B). This process of choosing initial experiments continues until the absolute difference between the maximum and the minimum yields/productivity is greater than a user-defined initial cutoff (5% MT yield was used for the valine dataset and 10 μg/gCDW/hr was used for the neurosporene dataset). These chosen initial experiments can be then labeled into two classes, “high” and “low”, based on their yield/productivity and the classifier is trained on these experiments and proposes subsequent experiments with predicted high chemical yield/productivity. The flowchart of the entire process is shown in Figure 3. The suggested subsequent experiment is the farthest or closest point on the “high” labeled side of the hyperplane, as certainty about the experimental outcome increases with distance from the decision boundary. After conducting the proposed experiment, the result is used to update the high/low cutoff used to classify all performed experiments (cutoff equals the average of the maximum and minimum yield/productivity across the previously selected experiments) and to train the next iteration’s SVM classifier. The SVM hyperplane might not change in each iteration as it depends on the support vectors. The process of suggesting experiments stops when there is no significant improvement in the yield (Figure 3C.i) or when no additional high yield/productivity experiments are predicted. Additionally, feature selection (Figure 3C.ii) can be performed by analyzing the weights of individual features. Classification using the MATLAB multinomial logistic regression function (mnrfit) was also performed on the valine dataset to identify the significance of each feature.

### Strains and plasmids

To evaluate how expression of different valine biosynthesis and exporter genes (Figure 1) impacts valine production, a derivative of *E. coli* strain PYR003 (BW25113 *aceE::kan ΔgdhA ΔpoxB NdhA)* with genotype BW25113 *ΔaceEΔgdhAΔpoxBΔldhAΔr•ecA* (PYR003a) was used as a background strain. PYR003 produces high yields of pyruvate from glucose and acetate (0.75 g pyruvate/g substrate) (X. Zhang and J.L. Reed, unpublished data). The valine biosynthesis genes *(ilvBN*DEIH*C/C*)* and valine exporter *(ygaZH (38))* genes were cloned onto two separate plasmids to allow combinatorial testing with varying expression levels. Valine production genes were either cloned from the *E. coli* K-12 MG1655 chromosome (in the case of *ilvBDEIC* and *ygaZH)* or were generated via overlap extension PCR (in the case of *ilvC*, ilvN\**, and *ilvH*)*. The *ilvC\** gene (containing mutations A71S, R76D, S78D, and Q110V and referred to previously as *ilvC*^6E6-his6^ *(35)*) prefers NADH instead of NADPH as a cofactor. The *ilvN\** gene (containing mutations G20D, V21D, and M22F and referred to previously as *ilvN*^mut^*(34))* and *ilvH\** gene (containing mutations G14D and S17F, referred to previously as *ilvH^G41A,C50T^(34))* are feedback-resistant mutants of *ilvN* and *ilvH*, respectively. The pTrc99A plasmid backbone *(39)* was used to express *ilvBN*DE*, while another plasmid backbone, pACYCtrc, was used to express *ilvC/C*, ilvIH\** and *ygaZH (40)*.

Multiple RBS sequences were used to generate different expression levels for the valine production genes (see Supplementary Table S1 for plasmid details). Specifically, RBS sequences were taken from either: 1) *de novo* designs from the RBS Calculator *(31)*; 2) published literature of characterized synthetic RBS sequences *(41)*; 3) chromosomal RBS sequences upstream of the gene’s genomic locus; or 4) RBS sequences already present on the plasmid backbones. RBS sequences generated by the RBS Calculator used the following input parameters: 1) Organism: *E. coli* K-12 MG1655; 2) free energy model v1.1; 3) 100 bp of the coding sequence; and 4) 20 bp upstream of the start codon.

### Media and culture conditions

All valine yield experiments were performed in 250 mL, baffled shake flasks containing 50 mL of MOPS-buffered minimal media *(42)* supplemented with 0.1 g/L sodium acetate, 2 g/L glucose, 100 μg/L thiamine hydrochloride, 100 mg/L of ampicillin, and 34 mg/L of chloramphenicol. Electro-competent PYR003a cells were prepared, double electroporated with two plasmid combinations, and incubated overnight at 37°C on Luria-Bertani broth *(43)* agar plates supplemented with 4 g/L glucose, 100 mg/L of ampicillin, and 34 mg/L of chloramphenicol. Subsequently, a minimum of two biological replicate colonies were picked for all experiments and sub-cultured in 10 mL of supplemented MOPS-buffered minimal media (as detailed above) for 24 hours at 37°C in a shaker at 225 RPM. Cells were then centrifuged, washed, and used to inoculate the 250 mL flasks to a starting OD_600_ of 0.01. Shake flasks were capped and wrapped with paraffin film to prevent evaporation and incubated for 48 hours. No isopropyl β-d-thiogalactopyranoside (IPTG) was added to the media, so transcription of the valine production genes from the plasmids was based on leaky expression

### Glucose and valine quantification

Prior to valine quantification, complete glucose utilization was verified for all experiments via an enzymatic assay (Glucose (GO) Assay Kit, Sigma-Aldrich) to ensure accurate yield calculations. Valine was quantified with a [1-^13^C]valine internal standard (Cambridge Isotope Laboratories) using an isotope-ratio method and gas chromatography-mass spectrometry (GC-MS) *(44)*. A known amount of a [1-^13^C]valine was added to samples containing unlabeled valine, dried at 90°C, and derivatized with *N*-*tert*-buty1-dimethylsilyl-*N*-methyltrifluoroacetamide plus 1%*tert*-butyl-dimethylchlorosilane at 90°C for 30 minutes to increase volatility and thermal stability required for GC-MS analysis. Samples were then run on a single quadrupole GC-MS QP2010S (Shimadzu) in electron ionization mode equipped with an Rtx-5ms (Restek) low-bleed, fused-silica column for separation with helium as a carrier gas operating under linear velocity control mode with a split ratio of 0.50 and a column flow of 1.50 mL/min. The temperature program for valine separation began with holding the oven temperature at 100°C for 5 minutes, ramping up at 25°C/min to 300°C, and holding for 5 minutes. Operating parameters included an injection temperature of 240°C, ion source temperature of 260°C, interface temperature of 240°C, and a mass scan range of 100-450 *m/z*. Then, an appropriate fragment *(45)* containing the labeled carbon from the internal standard was used to calculate the ^12^C/^13^C ratio and, subsequently, the concentration of the sample after correcting for isotopic impurity of the internal standard and for natural abundance of ^13^C using a freely available software, IsoCor *(46)*. This method was tested on samples with known concentrations of unlabeled valine ranging from 0.5 mM to 80 mM; predicted values were plotted against known values with a fit of y=0.9987x (with y=x being the most accurate). Measured valine yields were compared to the MT yield (0.644 g valine/g carbon source), the latter calculated from flux balance analysis *(47)* of the iJR904 *E. coli* genome-scale metabolic model *(48)* using the amounts of glucose (2 g/L) and acetate (0.072 g/L) present in the supplemented MOPS minimal medium.

## ACKNOWLEDGEMENTS

This work was funded in by the Office of Science (BER), U.S. Department of Energy (DE-SC0008103), the U.S. Department of Energy Great Lakes Bioenergy Research Center (DOE BER Office of Science DE-FC02-07ER64494 and DE-SC0018409), and by a grant from the Keck Foundation.

## Abbreviations

(LOOCV): leave-one-out cross-validation
(MT): maximum theoretical
(RBS): ribosome binding site
(SVM): Support Vector Machine
(UCB): upper confidence bound

## SUPPLEMENTARY INFORMATION

**Supplementary Figure S1:**
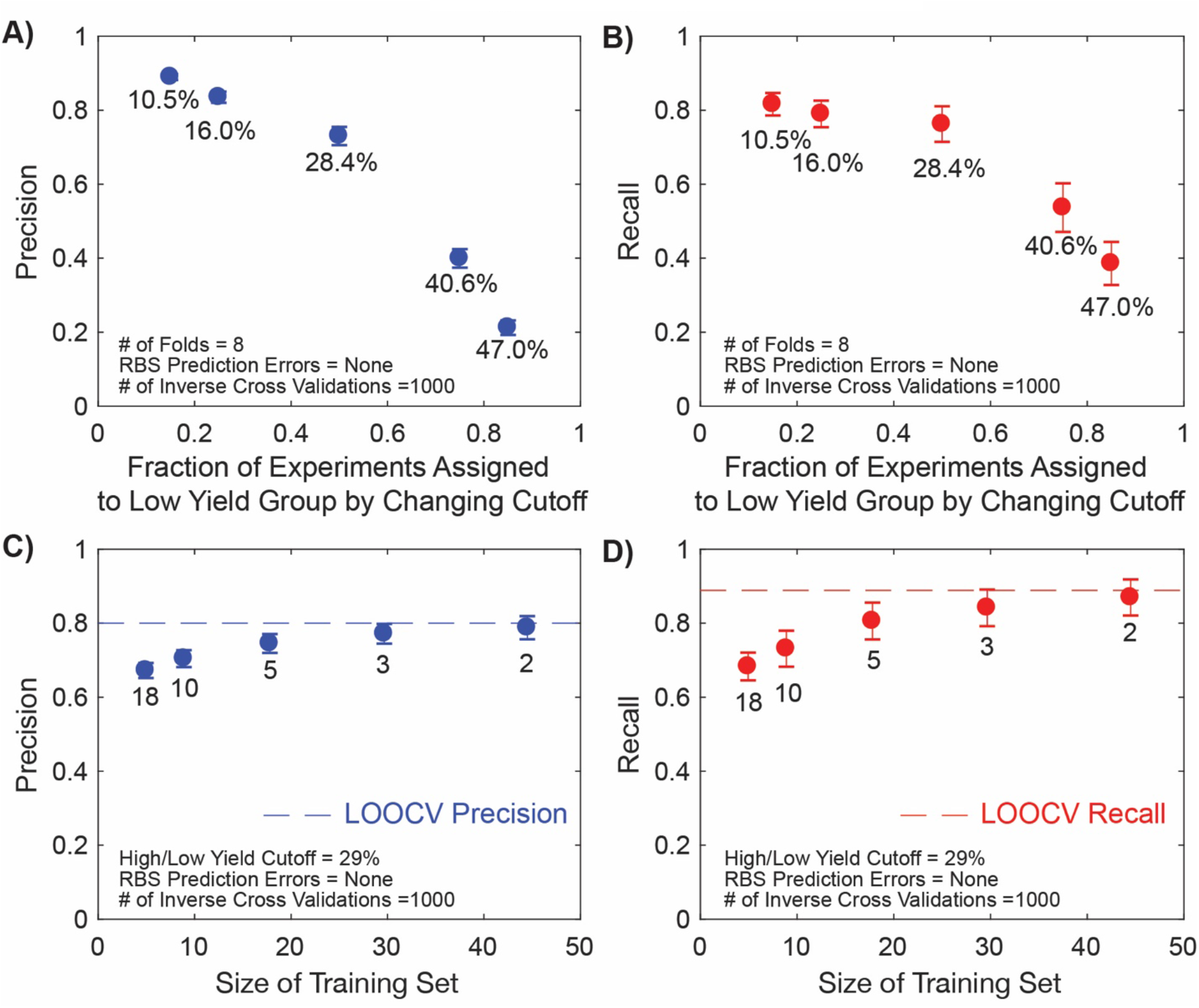
Sensitivity of the SVM Classifier to High/Low Cutoffs and Training Set Size. A cutoff was used to assign the valine experiments to one of two groups, either a high yield experiment or a low yield experiment. Panels (**A**) and (**B**) shows the sensitivity of the SVM classifier’s precision and recall, respectively, to the cutoff used to assign experiments to different groups. The cutoffs were varied so that 15%, 25%, 50%, 75%, or 85% of all the experiments were assigned to the low yield experiment group. Results in (**A**) and (**B** were generated by taking the average and standard deviation (error bars) of 1,000 inverse eight-fold cross-validations assuming no errors in the predicted RBS strengths. Panels (**C**) and (**D**) shows the sensitivity of the SVM classifier’s precision and recall, respectively, to the number of experiments included in the training dataset (by varying the number of folds used in the inverse crossvalidation). The number of folds were varied (18, 10, 5, 3, and 2) so that the size of the training datasets were around ~5, ~9, ~18, ~30, and ~45. Numbers below each point indicate cutoff (% MT Yield) used to generate each result. Results in (**C**) and (**D**) were generated by taking the average and standard deviation (error bars) of 1,000 inverse fold cross-validations using a yield cutoff of 29% MT yield (to assign experiments to separate groups) and assuming no errors in the predicted RBS strengths. The dashed lines in panels (**C**) and (**D**) show the precision and recall values from the LOOCV analysis (with a training set size of 90). Numbers below each point indicate number of folds used to generate each result.

**Supplementary Figure S2:**
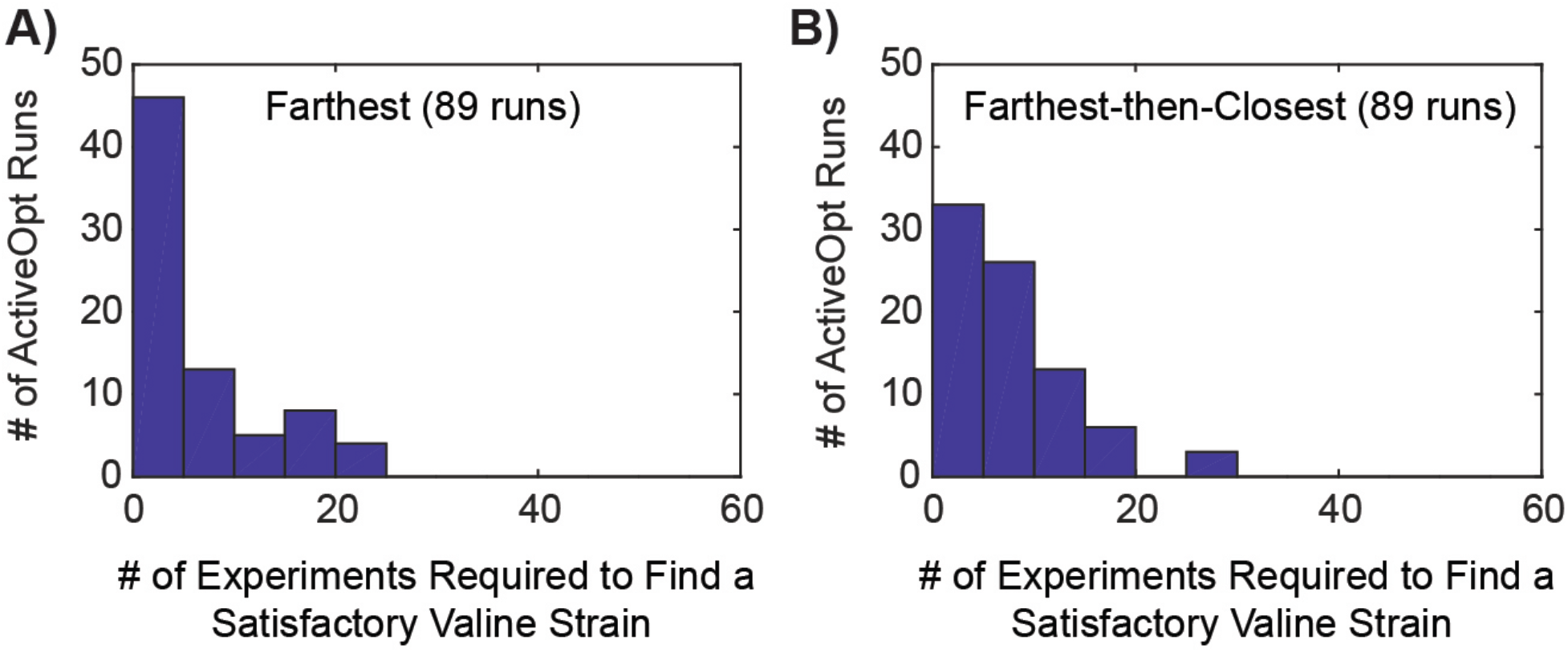
Number of Valine Experiments Needed to Find a Satisfactory Valine Strain. The figure shows histograms of the number of experiments needed to find a satisfactory valine strain for 89 different ActiveOpt runs (using each valine experiment as a first initial experiment) using either the farthest-from-the-hyperplane (Panel A) and farthest-then-closest-to-the-hyperplane (Panel B) approach. Panel A is the same as that shown in Figure 4C and is repeated for comparative purposes.

**Supplementary Figure S3:**
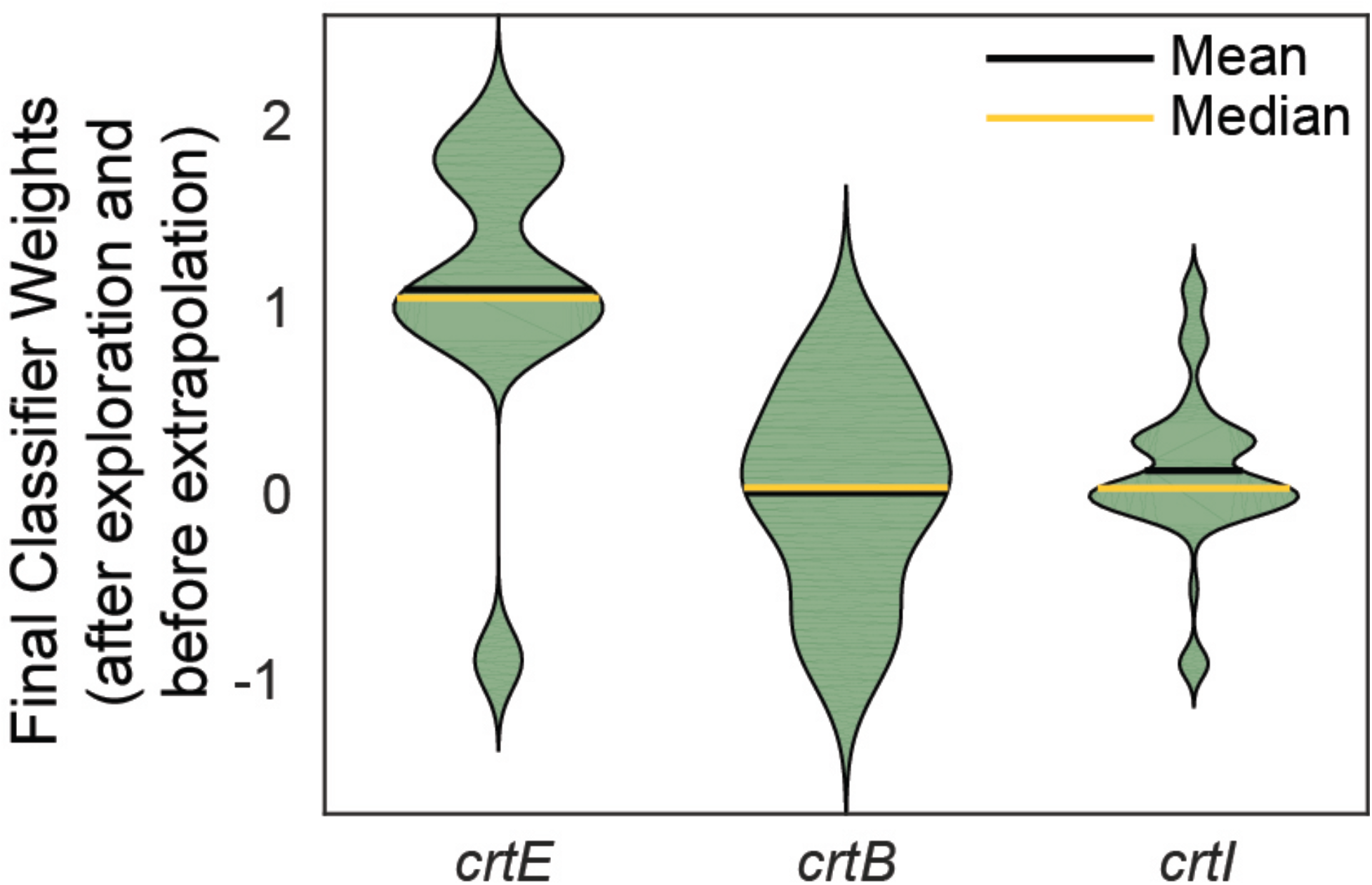
Distribution of Feature Weights for Final ActiveOpt Classifiers. This figure shows a violin plot (where the outer shape width is proportional to frequency of occurrence and the black and red bars indicates the mean and median values, respectively) for the distribution of weights for the three features (standardized RBS strengths for *crtE, crtB*, and *crtI*) across the final 73 ActiveOpt classifiers generated using the farthest-from-the-hyperplane approach. Each ActiveOpt run was generated using a different exploration experiment (the first 73 experiments reported by Farasat et al.) as a first initial experiment. The final classifier is when no more remaining experiments are predicted to be high yield.

**Supplementary Figure S4:**
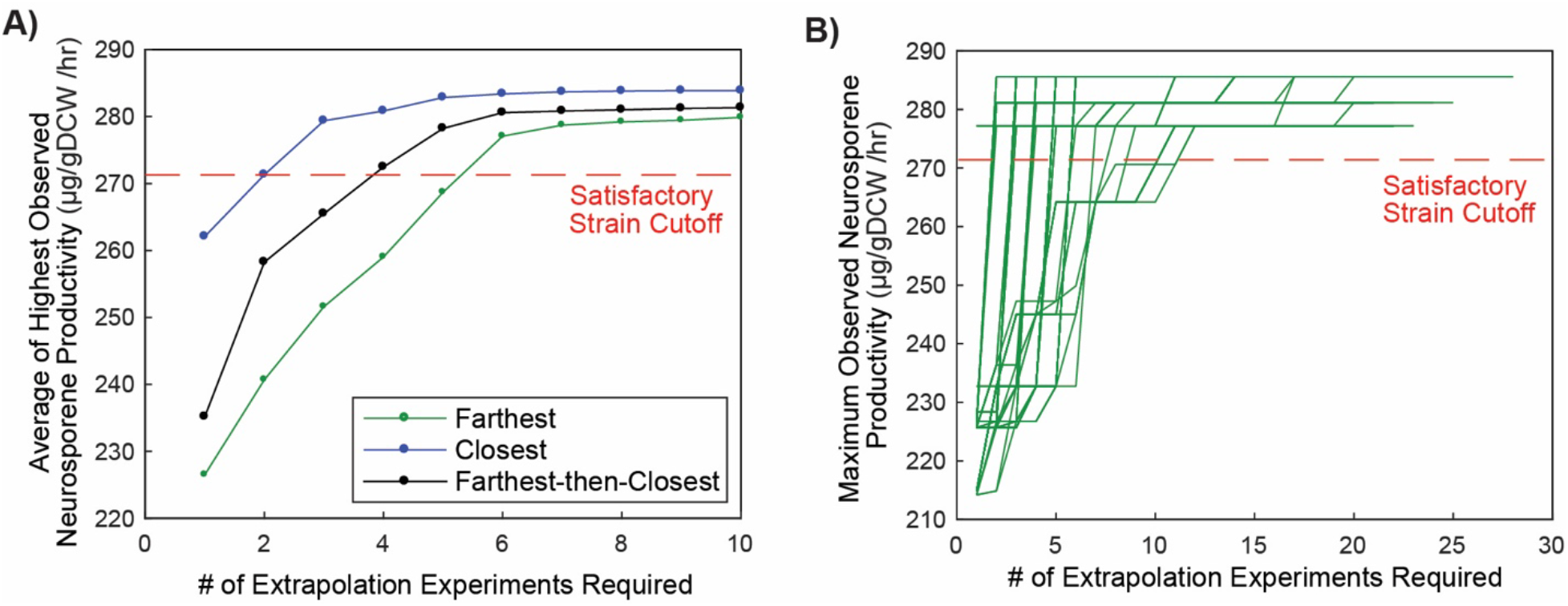
ActiveOpt Applied to Extrapolation Experiments from the Neurosporene Productivity Case Study. Panel (A) shows for different ActiveOpt next experiment selection approaches, the average from the 73 ActiveOpt runs of the highest observed neurosporene productivity as a function of the number of extrapolation experiments performed. The closest-to-the-hyperplane is shown in blue, the farthest-from-the-hyperplane is shown in green, and farthest-then-closest-to-the-hyperplane is shown in black. Panel (B) shows how the highest observed neurosporene productivity varies as a function of the number of extrapolation experiments performed using the farthest-from-the-hyperplane approach. Each of the 73 curves was generated by ActiveOpt starting from the final classifiers generated by ActiveOpt using the exploration experiments.

**Supplementary Table S1.** Plasmids and strains used in valine experiments. See Supplementary excel file.

**Supplementary Table S2.** Measured valine yields using different combinations of plasmids in PYR003a. See Supplementary excel file.

**Supplementary Table S3.** Performance of different techniques to select the next ActiveOpt experiment on the valine dataset.

|  | Random | Closest-to-the-Hyperplane | Farthest-from-the-Hyperplane | Farthest-then-Closest-to-the-Hyperplane |
| --- | --- | --- | --- | --- |
| # of ActiveOpt runs | 1000 | 89 | 89 | 89 |
| # of ActiveOpt runs that found a satisfactory strain<sup>a</sup> | 1000 | 83 | 76 | 81 |
| # of experiments to find a satisfactory strain<sup>b</sup> | 13 | 14 | 8 | 10 |
| # of ActiveOpt runs that found a satisfactory strain in <10 experiments<sup>c</sup> | 475 | 41 | 59 | 55 |
| average # of expts until no predicted high yield experiments remain<sup>d</sup> | NA<sup>g</sup> | 54.0 | 24.3 | 38.6 |
| Average precision<sup>e</sup> | NA<sup>g</sup> | 0.91 | 0.95 | 0.92 |
| Average recall<sup>f</sup> | NA<sup>g</sup> | 0.69 | 0.35 | 0.54 |

<sup>a</sup> Each run was started from a different first initial experiment
<sup>b</sup> The total number of experiments needed for the average (across all 89 runs) highest observed valine yield to exceed 95% of the measured maximum yield
<sup>c</sup> The number of runs that found a strain in less than 10 total experiments which had at least 95% of the measured maximum yield
<sup>d</sup> The average number of experiments suggested by ActiveOpt to be performed until no additional experiments are predicted to have high yield (i.e., the number of experiments needed to generate the final ActiveOpt classifiers)
<sup>e</sup> The average precision (across all 89 runs) for the final ActiveOpt classifiers when predictions were made for all 89 experiments
<sup>f</sup> The average recall (across all 89 runs) for the final ActiveOpt classifiers when predictions were made for all 89 experiments
<sup>g</sup> NA indicates not applicable since no classifier is generated using the random experiment selection approach.

**Supplementary Table S4.**
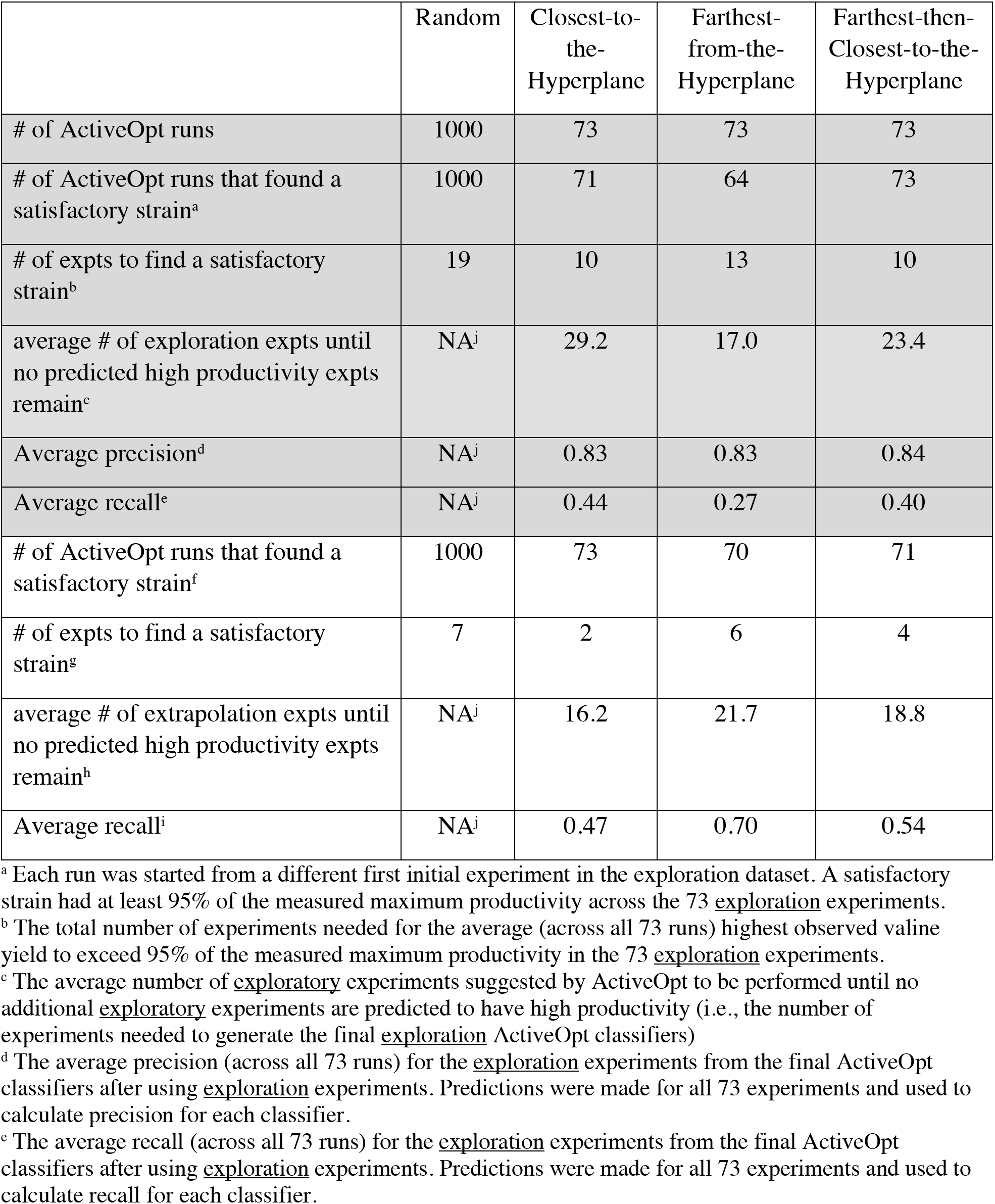
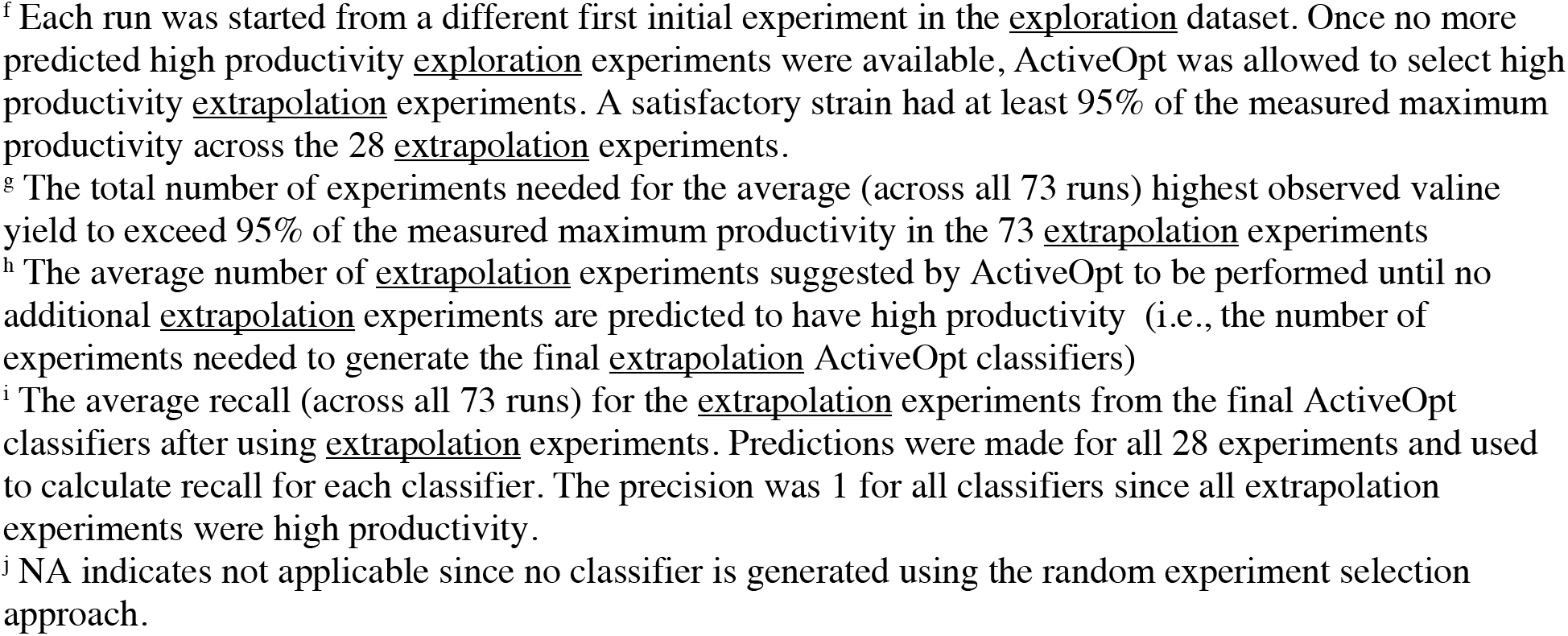
Performance of different techniques to select the next ActiveOpt experiment on the neurosporene dataset. In grey are results from the exploration experiments and in white the extrapolation experiments.

|  | Random | Closest-to-the-Hyperplane | Farthest-from-the-Hyperplane | Farthest-then-Closest-to-the-Hyperplane |
| --- | --- | --- | --- | --- |
| # of ActiveOpt runs | 1000 | 73 | 73 | 73 |
| # of ActiveOpt runs that found a satisfactory strain<sup>a</sup> | 1000 | 71 | 64 | 73 |
| # of expts to find a satisfactory strain<sup>b</sup> | 19 | 10 | 13 | 10 |
| average # of exploration expts until no predicted high productivity expts remain<sup>c</sup> | NA<sup>j</sup> | 29.2 | 17.0 | 23.4 |
| Average precision<sup>d</sup> | NA<sup>j</sup> | 0.83 | 0.83 | 0.84 |
| Average recall<sup>e</sup> | NA<sup>j</sup> | 0.44 | 0.27 | 0.40 |
| # of ActiveOpt runs that found a satisfactory strain<sup>f</sup> | 1000 | 73 | 70 | 71 |
| # of expts to find a satisfactory strain<sup>g</sup> | 7 | 2 | 6 | 4 |
| average # of extrapolation expts until no predicted high productivity expts remain<sup>h</sup> | NA<sup>j</sup> | 16.2 | 21.7 | 18.8 |
| Average recall<sup>i</sup> | NA<sup>j</sup> | 0.47 | 0.70 | 0.54 |

<sup>a</sup> Each run was started from a different first initial experiment in the exploration dataset. A satisfactory strain had at least 95% of the measured maximum productivity across the 73 exploration experiments.
<sup>b</sup> The total number of experiments needed for the average (across all 73 runs) highest observed valine yield to exceed 95% of the measured maximum productivity in the 73 exploration experiments.
<sup>c</sup> The average number of exploratory experiments suggested by ActiveOpt to be performed until no additional exploratory experiments are predicted to have high productivity (i.e., the number of experiments needed to generate the final exploration ActiveOpt classifiers)
<sup>d</sup> The average precision (across all 73 runs) for the exploration experiments from the final ActiveOpt classifiers after using exploration experiments. Predictions were made for all 73 experiments and used to calculate precision for each classifier.
<sup>e</sup> The average recall (across all 73 runs) for the exploration experiments from the final ActiveOpt classifiers after using exploration experiments. Predictions were made for all 73 experiments and used to calculate recall for each classifier.
<sup>f</sup> Each run was started from a different first initial experiment in the exploration dataset. Once no more predicted high productivity exploration experiments were available, ActiveOpt was allowed to select high productivity extrapolation experiments. A satisfactory strain had at least 95% of the measured maximum productivity across the 28 extrapolation experiments.
<sup>g</sup> The total number of experiments needed for the average (across all 73 runs) highest observed valine yield to exceed 95% of the measured maximum productivity in the 73 extrapolation experiments
<sup>h</sup> The average number of extrapolation experiments suggested by ActiveOpt to be performed until no additional extrapolation experiments are predicted to have high productivity (i.e., the number of experiments needed to generate the final extrapolation ActiveOpt classifiers)
<sup>i</sup> The average recall (across all 73 runs) for the extrapolation experiments from the final ActiveOpt classifiers after using extrapolation experiments. Predictions were made for all 28 experiments and used to calculate recall for each classifier. The precision was 1 for all classifiers since all extrapolation experiments were high productivity.
<sup>j</sup> NA indicates not applicable since no classifier is generated using the random experiment selection approach.

**Supplementary Table S5.** Average weights across final ActiveOpt classifiers generated from the 73 neurosporene exploration experiments, with each experiment chosen as a first initial experiment.

| **Next Experiment Selection Approach** | *crtE* | *crtB* | *crtI* |
| --- | --- | --- | --- |
| closest-to-the-hyperplane | 0.98 | -0.04 | 0.11 |
| farthest-from-the-hyperplane | 1.07 | -0.03 | 0.09 |
| farthest-then-closest-to-the-hyperplane | 0.96 | 0.03 | 0.07 |

